# Neurogrid simulates cortical cell-types, active dendrites, and top-down attention

**DOI:** 10.1101/2021.05.14.444265

**Authors:** Ben Varkey Benjamin, Nicholas A Steinmetz, Nick N Oza, Jose J Aguayo, Kwabena Boahen

**Affiliations:** Department of Electrical Engineering, Stanford University, Stanford, CA 94305, USA; Department of Neurobiology, Stanford University, Stanford, CA 94305, USA; Department of Bioengineering, Stanford University, Stanford, CA 94305, USA

**Author notes:** The data that support the findings of this study are available from the corresponding author upon reasonable request. Correspondence and requests should be addressed to K.B.

## Abstract

A central challenge for systems neuroscience and artificial intelligence is to understand how cognitive behaviors arise from large, highly interconnected networks of neurons. Digital simulation is linking cognitive behavior to neural activity to bridge this gap in our understanding at great expense in time and electricity. A hybrid analog-digital approach, whereby slow analog circuits, operating in parallel, emulate graded integration of synaptic currents by dendrites while a fast digital bus, operating serially, emulates all-or-none transmission of action potentials by axons, may improve simulation efficacy. Due to the latter’s serial operation, this approach has not scaled beyond millions of synaptic connections (per bus). This limit was broken by following design principles the neocortex uses to minimize its wiring. The resulting hybrid analog-digital platform, Neurogrid, scales to billions of synaptic connections, between up to a million neurons, and simulates cortical models in real-time using a few watts of electricity. Here, we demonstrate that Neurogrid simulates cortical models spanning five levels of experimental investigation: biophysical, dendritic, neuronal, columnar, and area. Bridging these five levels with Neurogrid revealed a novel way active dendrites could mediate top-down attention.

## 1 MULTISCALE NEURAL SIMULATIONS

A central challenge for systems neuroscience and artificial intelligence is to understand how cognitive behaviors arise from large, highly interconnected networks of neurons^1^. Bridging this gap in our understanding calls for large-scale yet biophysically realistic neural simulations to span levels of experimental investigation (biophysical, dendritic, neuronal, columnar, and area)^2^. Digital simulation bridges these levels, but the expense in time and electricity is great^3^.

A Titan RTX GPU card consumes electricity at a rate of 250W to simulate a cortex model with 4 million neurons connected by 24 billion synapses at a speed 510-fold slower than real-time (i.e. each biological second takes 510 sec)^4^. These simulations are challenging because each neuron *distributes* its output to thousands of neurons and, in turn, *aggregates* inputs from thousands of neurons.

In the digital paradigm, replicating signals for distribution as well as summing signals for aggregation occurs entirely within computing cores (facilitated by programming constructs such as *MapReduce*). The communication network interconnecting these cores simply routes messages by relaying them across links that connect a core to its immediate neighbors. Sharing these core-to-core wires instead routing dedicated point-to-point wires, as the cortex does, ameliorates the difficulty of mapping a threedimensional brain onto a two-dimensional chip^5^.

With shared wires communicating addresses that signal arrival of an action potential, or spike, at particular synapse, traffic scales as the total number of synaptic connections times the presynaptic population’s average spike-rate^6^. This *address-event* bus has been successfully used to build networks with thousands of neurons with a few hundred synaptic connections each^7^. It has not scaled beyond millions of synaptic connections, the point at which traffic saturates the bus’ signaling rate.

To break this communication bottleneck, Neurogrid adopts a hybrid analog-digital approach that follows design principles the neocortex uses to minimize its wiring^8^. Neurogrid uses fast digital routers, operating serially, to replicate signals for distribution and uses slow analog circuits, operating in parallel, to sum signals for aggregation. It follows the neocortex’s design principles by performing both distribution and aggregation hierarchically. This neuromorphic architecture scales to billions of synaptic connections, between up to a million neurons.

Here, we demonstrate that Neurogrid simulates cortical models spanning five levels of experimental investigation: biophysical, dendritic, neuronal, columnar, and area (Sec. 4 thru Sec. 6). Bridging these five levels with Neurogrid revealed a novel way active dendrites could mediate top-down attention. We begin with a brief review of Neurogrid’s neuromorphic architecture (Sec. 2) and simulation environment (Sec. 3); detailed descriptions are available elsewhere^8^.

## 2 HIERARCHICAL DISTRIBUTION & AGGREGATION

A neuron’s axonal arbor distributes signals by hierarchically replicating them at branch points. Placing an axon’s branch points as close as possible to its terminals replaces multiple axon segments with a single segment^9^ (Fig. 1a, *top*). This principle minimizes the axonal arbor’s wiring.

**Figure 1.**
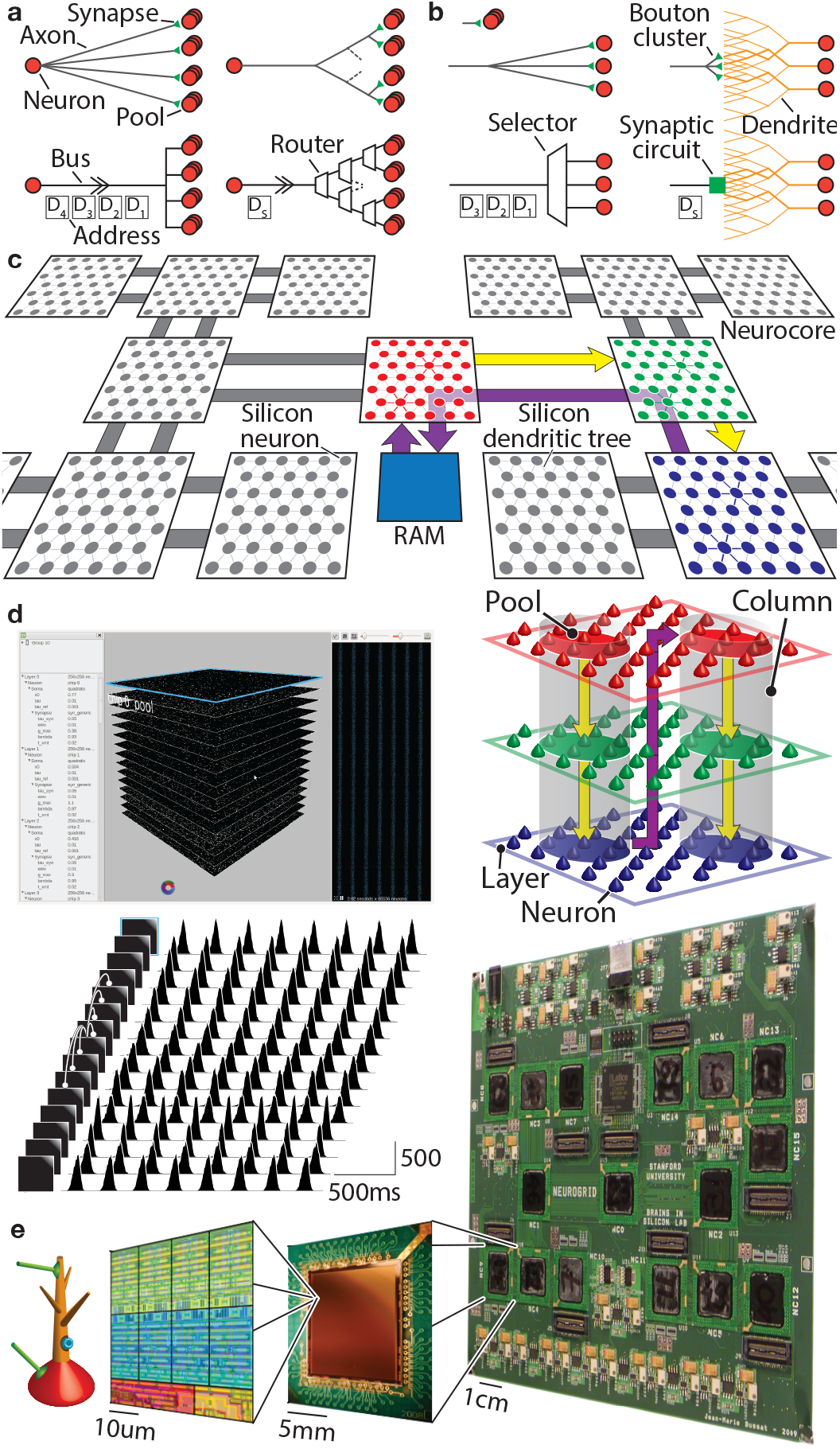
Modeling the cortex on Neurogrid. **a**, Hierarchical distribution and **b**, hierarchical aggregation in neural (*top*) and neuromorphic (*bottom*) networks: The traffic on a digital bus that emulates spike distribution by an axonal arbor is reduced by mimicking axonal and dendritic branching patterns. **c**, Mapping cortical columns: Cell layers (red, green and blue), intercolumn projections (purple), and intracolumn collaterals (yellow) are mapped onto a different Neurocores, off-chip RAM (on a daughterboard), and on-chip RAM (in each Neurocore), respectively. **d**, Simulating 1M neurons with 8G synaptic connections in realtime. User-interface (top, *left* panel to *right* panel): Displays model parameters, activity in all layers, and spike trains from a selected layer (the first one). Layer connectivity and spike histograms (*bottom*, *left* & *right*): Each layer inhibits itself and its three immediate neighbors on either side (only the eighth layer’s connectivity is shown). These local interactions synchronize activity globally. **e**, Hardware (*left* to *right*). Silicon neuron schematic and layout: Distinct soma and dendrite compartments express spike-generating, ligand-gated, and voltage-gated conductances. The last two occupy the most area (four types each). Neurocore chip: Has 65,536 silicon neurons, tiled in a 256 × 256 array, and routing circuitry for interchip communication. Neurogrid board: Has 16 Neurocores, connected in a binary-tree network (as in **c**).

Moreover, extending several dendritic branches to meet a terminal branch of an axon allows that branch to make multiple synapses (bouton cluster)^10^ (Fig. 1b, *top*). Synaptic signals from many axons sum in a dendritic branch and these branches’ signals aggregate hierarchically. This principle minimizes the dendritic tree’s wiring^9^.

Hierarchical distribution is emulated by interconnecting silicon-neuron arrays with a tree-like (rather than a mesh-like) network^11,12^. That allows address-events to be replicated close to their destination and reduces traffic on the array-to-array links (Fig. 1a, *bottom*). Emulating an axon terminal’s bouton cluster allows a single delivered address-event to evoke postsynaptic potentials in several silicon neurons (Fig. 1b, *bottom*). This reduces traffic further.

Hierarchical aggregation is emulated efficiently by modeling multiple overlapping dendritic trees with a single two-dimensional resistive grid. This shared analog circuit replicates the exponential spatial decay that transforms postsynaptic potentials into current delivered to a dendritic tree’s trunk^13^.

## 3 NEUROGRID

Emulating axonal arbors and dendritic trees enables Neurogrid to simulate cortical models scalably and efficiently. A cortical area is modeled by a group of *Neurocores* (Fig. 1c); Neurogrid has sixteen of these chips. Each cell layer (or cell type) is mapped onto a different Neurocore’s two-dimensional silicon neuron array. Circular pools of neurons centered at the same (*x*, *y*) location on these Neurocores model a cortical column^15^.

Intercolumn axonal projections are routed by using the presynaptic neuron’s address to retrieve the target columns’ centers (*x, y*) from an off-chip random-access memory (first distribution level). This memory is programmed to replicate the neocortex’s function-specific intercolumn connectivity^16,17^.

Intracolumn axon collaterals are routed by copying the address-event to all of a cortical area’s Neurocores using the interchip tree network; unneeded copies are filtered using an on-chip memory (second distribution level). This memory is programmed to replicate the neocortex’s stereotyped intracolumn connectivity^18^.

Intralayer dendritic trees—arborizing over a circular disc centered on the cell body—are realized using the resistive grid mentioned earlier. Its space constant is adjusted electronically to match the arbor’s radius; a transistor-based implementation makes this possible^19^.

With intercolumn projections, intracolumn collaterals, and intralayer dendrites, five thousand synaptic connections may be made by: (1) Routing an address-event to ten columns; (2) copying it to six layers in each column; and (3) evoking postsynaptic potentials in a hundred neighboring neurons in all but one layer (a 5.6-neuron arbor radius).

To demonstrate these routing mechanisms’ functionality and scalability, we used Neurogrid to simulate a recurrent network of a million interneurons with billions of inhibitory synaptic connections (Fig. 1d). The interneurons were organized in sixteen 256×256-neuron arrays that formed a torus; the first and last layers were neighbors. Intracolumn collaterals routed an interneuron’s spike to the three nearest layers on each side as well as back to its own layer. The inhibition evoked decreased exponentially with distance, due to spatial decay in the intralayer dendritic trees. Half of the inhibition a soma received came from 8,000 neurons in a cylinder centered on it, 7 layers in height and 19 neurons in radius.

These local inhibitory interactions synchronized the interneurons globally, reproducing previous findings^20^. Spiking activity waxed and waned periodically (3.7 Hz), with individual interneurons skipping several cycles (0.42 spikes/s mean). Nevertheless, their spikes were entrained to the global rhythm across all 16 layers, demonstrating the functionality and scalability of Neurogrid’s routing mechanisms.

Software infrastructure facilitates performing large-scale neural simulations on Neurogrid. Firmware and drivers initialize and configure the hardware. Calibration procedures establish correspondence between parameters of neuronal models and their silicon analogs^21,22^. A mapping tool translates multiscale model descriptions (written in Python) into hardware configurations (programmed in memory). A user interface interacts with simulations in real-time. And visualization widgets render activity or plot spikes of up to a million neurons in real-time. For the details of this software infrastructure, see Methods.

## 4 NEOCORTICAL CELL-TYPES

Neurogrid’s neuron model has separate soma and dendrite compartments (Fig. 1e). The soma compartment expresses conductances that generate spikes and adapt their rate. The dendrite compartment expresses ligand and voltage-gated conductances (up to four types each). The programmed maximum synaptic or membrane conductance is scaled by a “gating variable”^23^. A pair of gating variables may be used to model a population of ion-channels that are activated *and* inactivated by voltage (e.g., T-type Ca channels), activated by ligand *and* voltage (e.g., NMDA receptors), or modulated by a second ligand (e.g., dopamine). For these components’ equations and parameters, see Methods.

To demonstrate the expressiveness of this neuron model, we simulated four prominent neocortical cell-types classified by Nowak et. al.^24^, namely fast spiking (FS), regular spiking (RS), intrinsic bursting (IB) and chattering (CH). FS neurons, normally associated with inhibitory interneurons, respond to a depolarizing current input with a high-frequency spike-train, showing little or no adaptation. RS neurons, generally associated with excitatory stellate cells, respond to a depolarizing current input with a low-frequency spike-train, showing adaptation. For both types of neurons, increasing the injected current increases the spike rate.

We modeled FS and RS neurons with the soma compartment alone, including adaptation for the latter (Fig. 2a,b). These models matched the FS neuron’s high spike-rate and relatively constant inter-spike intervals; the RS neuron’s low spike-rate and increasing inter-spike intervals; and both cell types’ overall spike-rate increase in with injected current.

**Figure 2.**
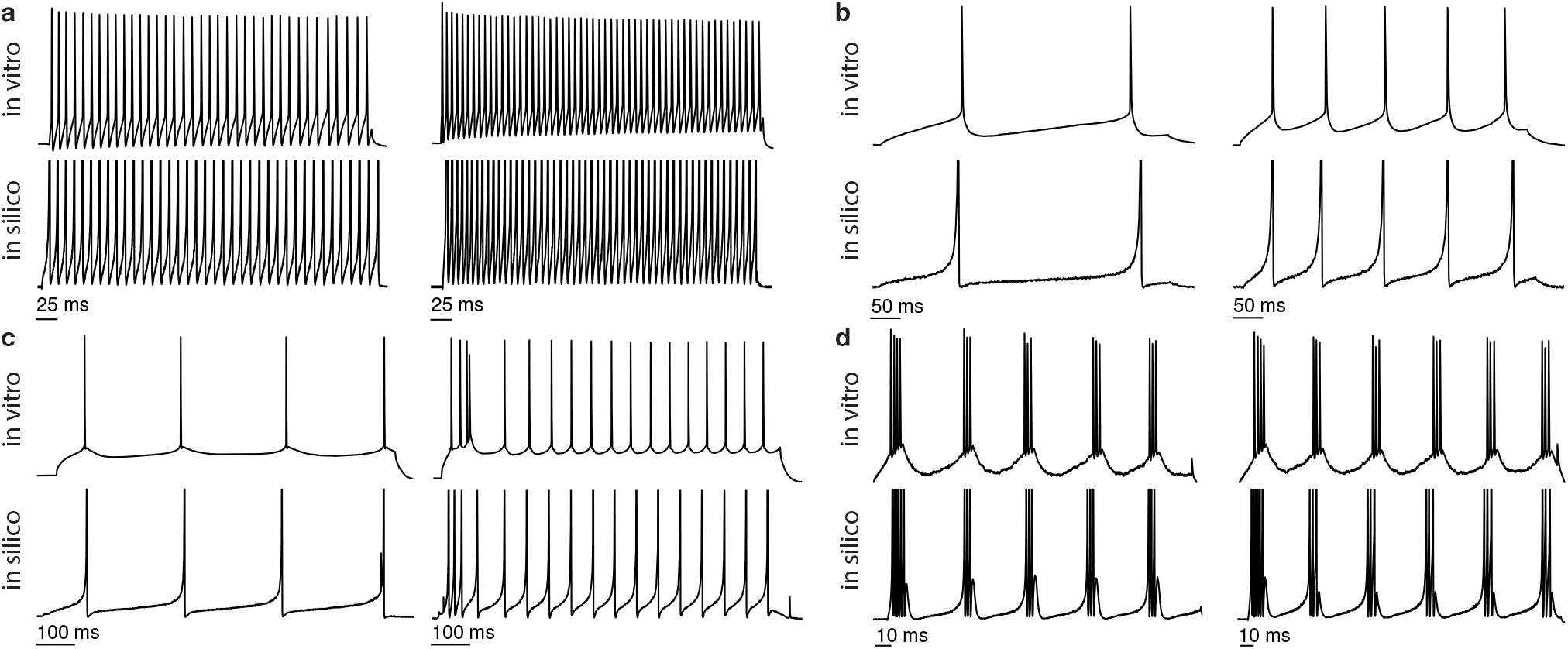
Different types of neocortical neurons and their Neurogrid simulations. Each quadrant shows membrane potentials recorded *in vitro* (upper traces) and *in silico* (lower traces) with two current injection levels (increasing from left to right). **a**, FS neuron. **b**, RS neuron. **c**, IB neuron. **d**, CH neuron. Over the ranges of input currents used (see Methods), the model’s spike-times closely match the biological neuron’s, even though the former are always reset to the same potential (not programmable). *In vitro* data adapted from Izhikevich^14^.

IB and CH neurons display more complex responses. IB neurons, generally associated with pyramidal cells, respond to a sufficiently large increase in injected current with a burst followed by single spikes, as observed in guinea-pig cingulate neocortex^25^. Bursting does not occur if the current is injected when the membrane is depolarized. CH neurons, also associated with pyramidal cells, exhibit fast rhythmic bursting, as observed in cat striate cortex^26^. Their burst frequency does not increase much with increasing injected current.

We modeled IB and CH neurons with the soma and dendrite compartments. Slow calcium channels that are deinactivated near the resting potential contribute to bursting^27^. Hence, we equipped the dendrite compartment with an activating as well as an inactivating gating variable (Fig. 2c). This model matched the onset of bursting in an IB neuron and the increase in tonic spike-rate with increasing injected current.

Rhythmic bursting could arise from bidirectional interactions between the soma and the dendrite^28^. Neurogrid’s neuron model supports this by allowing spikes to back-propagate from the soma compartment to the dendrite compartment (purely passive in this case). In turn, the dendrite depolarizes the soma. These reciprocal interactions produce repetitive spiking^29^. A build-up of K^+^ conductance with each spike terminates the burst; this conductance’s decay with time determines the inter-burst interval. This model matched the high spike-rate within a burst, the number of spikes per burst, and the moderate decrease in inter-burst interval with increasing injected current (Fig. 2d).

## 5 ACTIVE DENDRITES

Several lines of experimental evidence point to a critical role of active dendritic properties in sensorimotor processing. NMDA spikes in basal dendrites^30^ have been shown to sharpen sensory tuning of neocortical neurons^32^. Such spikes double an EPSP’s amplitude when they occur in basal dendrites of neocortical layer 5 pyramidal neurons. Coincidence of EPSPs (pre-spikes) and back-propagating action potentials (post-spikes) in apical dendrites^31^ has been shown to amplify correlated sensory and motor signals during active sensation^33^. Thus, active dendrites enhance sensorimotor processing.

To demonstrate that Neurogrid’s neuron model could express active dendritic behavior, we replicated the NMDA spike’s all-or-none behavior and the backpropagating spike’s amplifying action. A NMDA spike occurs when above some threshold a small increment of synaptic strength more than doubles the amplitude of the excitatory post-synaptic potential (EPSP), which then remains unchanged for further increments^30^. To replicate the NMDA spike, we expressed a NMDA synaptic conductance in a dendrite compartment as well as an AMPA conductance. When the AMPA conductance drove the dendrite’s potential above the NMDA conductance’s voltage-activation threshold, the postsynaptic potential doubled in amplitude, replcating the all-or-none behavior of an NMDA spike (Fig. 3a).

**Figure 3.**
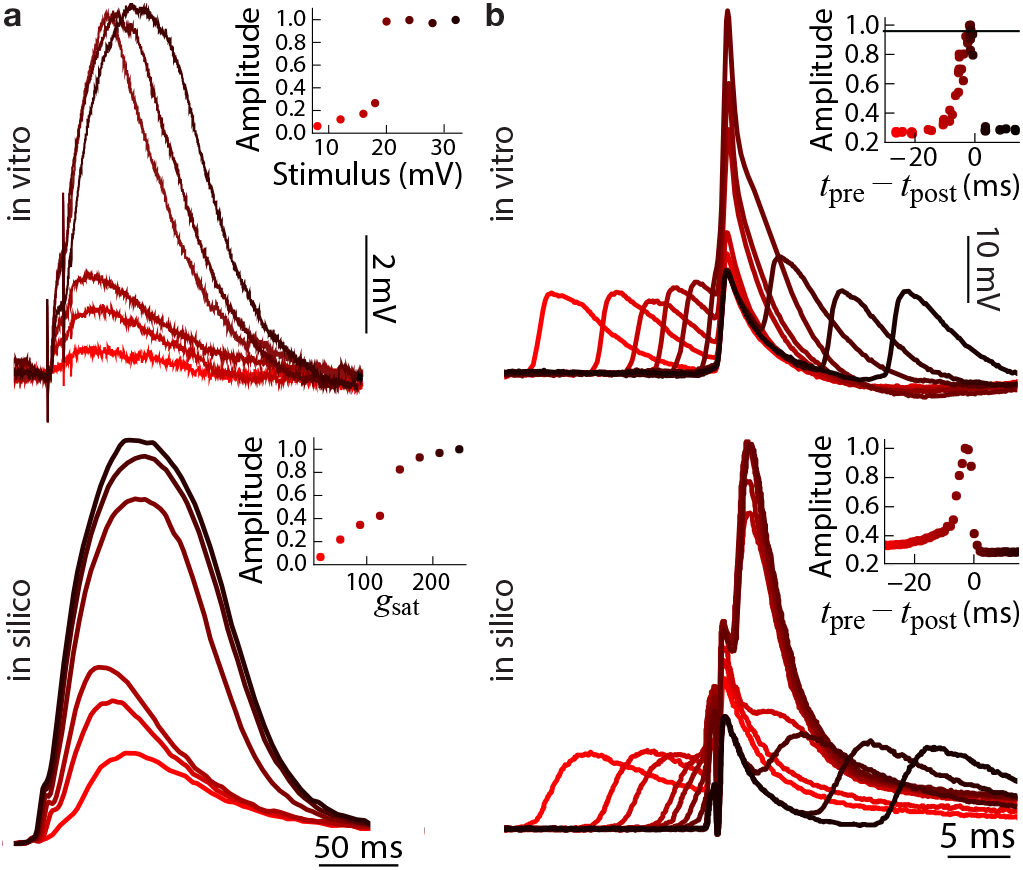
Active dendritic behaviors and their Neurogrid simulations. Each column compares *in vitro* (top) and *in silico* (bottom) membrane potential traces; insets plot traces’ peak values versus stimulus conditions. **a**, NMDA spikes in basal dendrites: The synaptic conductance was varied by stimulating afferent axons electrically *in vitro* or increasing the maximum synaptic conductance *in silico* (*g*_sat_). **b**, Coincidence detection in apical dendrites: The relative timing between the backpropagating spike (*t*_pre_) and the EPSP (*t*_post_) was varied. *In vitro* data in **a** and **b** adapted from Schiller et al.^30^ and Stuart and Hausser^31^, respectively.

To replicate the amplifying action of a backpropagating spike, we expressed sodium channels that activate and inactivate in a dendrite compartment as well as a synaptic conductance with timeconstant between NMDA and AMPA conductances’ to mimic a mix. When a backpropagating spike coincided with a postsynaptic potential, the sodium conductance activated and increased the dendrite’s voltage further. The pre-spike’s timing relative to the post-spikes’ when voltage peaked as well as the voltage’s asymmetric increase and decrease matched experimental observations in rat layer 5 pyramidal neurons^31^ (Fig. 3b). Replicating these *in-vitro* results did not require NMDA synapses’ voltage-dependence, consistent with previous models^31^.

## 6 TOP-DOWN ATTENTION

To demonstrate that Neurogrid can help relate cognitive phenomena to biophysical mechanisms, an important goal of neuroscience, we replicated the effects of spatially selective top-down attention on the responses of visual cortical neurons recorded in awake, behaving macaques^34^. Amplification of postsynaptic potentials by NMDA or sodium conductances in apical dendrites of pyramidal cells could contribute to this multiplicative interaction. The former may underlie spike-rate changes associated with feedback-driven figure-ground modulations in Macaque primary visual cortex^35^. This evidence led us to hypothesize that NMDA receptors could also account for the multiplicative changes in spike rate observed in visual cortical neurons during spatially selective attention^34^.

Unlike most computational models, Neurogrid models are subjected to heterogeneity. Once calibrated, the median parameter value across a Neurocore’s population of silicon neurons matches the specified model parameter value. Whereas its variance is determined by the fabrication process, resulting in a spike-rate distribution that spans a decade. For comparison, firing rates vary by over three decades across a population of cortical neurons^36^. Hence, showing that a proposed mechanism is robust to heterogeneity increases its biological plausibility.

We tested robustness of NMDA-mediated multiplicative gain to heterogeneity by simulating top-down and bottom-up interactions between a visual cortical area (V4) and a frontal cortical area (the Frontal Eye Field, FEF) on Neurogrid (Fig. 4a). V4 was mod-eled with an excitatory neuronal population (128 × 128) while FEF was modeled with an excitatory and an inhibitory population (both 128 × 128). Besides V4’s excitatory population, which had dendrite compartments, all others had just soma compartments.

**Figure 4.**
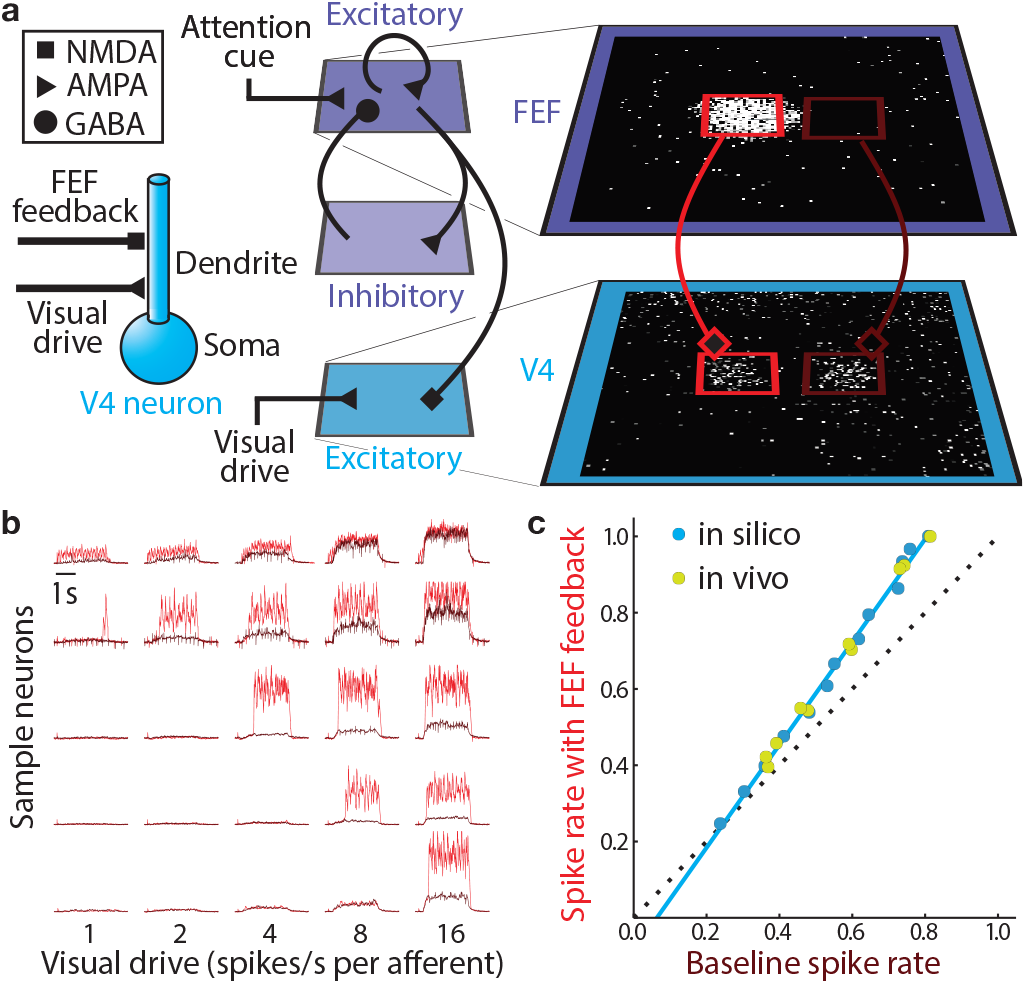
Modeling attention-related modulation of visual cortical responses. **a,** FEF-V4 model: Cued activity in FEF (*right*, white pixels within top 19 × 19 red rectangle) drives NMDA receptors in a patch of excitatory neurons at the corresponding location in V4 (*right*, white pixels within bottom red rectangle); neurons there and elsewhere (*right*, bottom brown rectangle) receive identical external visual drive. **b,** Heterogeneous responses in model V4: Dendritic potentials with and without FEF drive (red and brown, respectively) for five sample neurons (rows) sorted according to the visual drive required to cross NMDA synapse’s activation threshold. **c,** Population activity in model V4: The average spike-rate scales multiplicatively with NMDA-mediated FEF feedback (calculated for 361 neurons), matching the behavior of V4 neurons recorded from awake, behaving Macaques (adapted from McAdams and Maunsell^34^).

FEF activity, temporally sustained by local recurrent excitation and spatially constrained by local recurrent inhibition, represented the locus of attention in a 2D map of visual space, inspired by Ardid et al.’s (1D) attention model^37^. To explore NMDA receptors’ role in attentional modulation, unlike in the previous model, columnar feedback projections from FEF to V4 modulated activity of V4 neurons exclusively through NMDA synapses.

We discovered that heterogeneity in AMPA conductances realizes gradual gain modulation from abrupt NMDA threshold crossings. This heterogeneity (CV of 26%) caused dendrite compartments to cross NMDA’s voltage-activation threshold at different visual drives (Fig. 4b). This distributed threshold-crossing increased the V4 population’s spike-rate gradually with visual drive; it would otherwise have increased abruptly. This increase was steeper with FEF feedback, matching multiplicative changes with attention observed in macaque V4^34^ (Fig. 4c). This novel NMDA-mediated gain mechanism realizes multiplicative attentional modulation in a biological plausible manner.

## 7 CONCLUSIONS

Conductance-based synapses, active conductances, multiple dendritic compartments, spike back-propagation, and cortical cell types have been emulated in neuromorphic chips^29,38–45^. Our focus here was on the next step: Deploying these *in silico* neuronal components in multiscale modeling. We simulated cortical models with up to 1M neurons and 8G synaptic connections using 1800-fold less energy per synaptic activation than a GPU^4,8^ (120pJ versus 210nJ)^i^.

A hybrid analog-digital architecture made this possible. It implements hierarchical distribution and aggregation and maps cortical circuitry columnarly. Following these principles the neocortex uses to minimize its wiring reduced traffic between cores greatly. This neuromorphic approach enabled Neurogrid to simulate multiscale neural models that integrate findings across five levels of experimental investigation energy- and time-efficiently. Importantly, Neurogrid achieves scale and efficiency without sacrificing biophysical detail, and thus truly supports multiscale modeling.

Neurogrid’s neuronal model is sufficiently detailed to simulate various cortical cell types as well as active dendritic behavior and neuromodulatory effects. This combination of scale and detail enabled us to discover a novel gain mechanism for attentional modulation. This advance in simulation capacity could not only lead to a better understanding of neurological disorders such as ADHD, it could also help reverse-engineer the brain’s hybrid analog-digital computational paradigm, which could lead to artificial intelligence systems with billions of units and trillions of parameters that consume watts instead of kilowatts.

## Acknowledgements

We thank E. Kauderer-Abrams for calibration algorithm development; B. Softky and H. S. Seung for assistance in editing the manuscript; and K. Chin for administrative support. This work was supported by an NIH Director’s Pioneer Award (DPI-OD000965) and a NIH/NINDS Transformative Research Award (R01NS076460).

## Author Contributions

B.V.B. developed the synchrony, NMDA-spike, coincidence detection, and cortical neuron models. N.A.S. developed the attention model. N.N.O. and J.J.A. developed calibration, mapping, interaction and visualization software. K.B. directed the research effort. B.V.B., N.A.S. and K.B. wrote the manuscript.

## Author Information

K.B. is a co-founder and equity owner of Femtosense Inc. The remaining authors declare that they have no competing financial interests.

## METHODS

### Dimensionless neuron models

Neurogrid’s models of soma, dendrite, gating variables and synapses are dimensionless. A leaky integrate-and-fire model of a neuron^1^ is described by

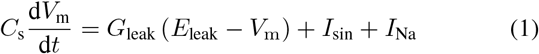

where *V*_m_ is the soma membrane potential, *C*_s_ is the soma membrane capacitance, *G*_leak_ is the soma leak conductance and *E*_leak_ its reversal potential, *I*_sin_ is the injected current and *I*_Na_ is the voltage-dependent sodium current that generates spikes. Rewriting the above equation with *E*_leak_ as the reference voltage:

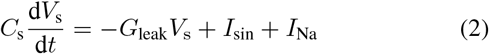

where *V*_s_ = *V*_m_ – *E*_leak_. We modeled *I*_Na_ (*V*_s_) as

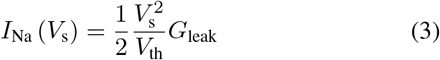

where *V*_th_ is the threshold voltage. By definition, it is the voltage at which the sodium conductance (d*I*_Na_/d*V*_s_) is equal to the leak conductance. At this voltage, the membrane potential stops decelerating and starts accelerating (inflection point), the onset of spiking. Substituting equation (3) into equation (2) and dividing by *G*_leak_*V*_th_ gives the dimensionless model for a Quadratic Leaky Integrate-and-Fire (QLIF) neuron^2,3^:

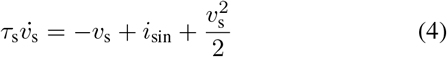

where *τ*_s_ = *C*_s_/*g*_leak_ (membrane time constant), *v*_s_ = *V*_s_/*V*_th_ and *i*_sin_ = *I*_sin_/ (*g*_leak_/*V*_th_). Thus, models of the form equation (1) can be expressed as dimensionless models of the form equation (4) by changing the reference voltage to *E*_leak_ and normalizing all voltages, conductances and currents by *V*_th_, *G*_leak_ and *G*_leak_*V*_th_, respectively.

The soma compartment’s dimensionless model is

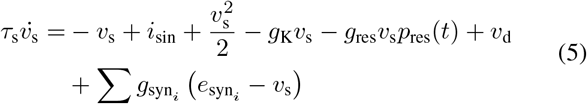

where *v*_s_ is the membrane potential, *τ*_s_ the membrane time constant, *i*_sin_ the input current, and *g*_syn_i__ and *e*_syn_i__ the conductance and reversal potential, respectively, of synapse *i*. *g*_res_ is activated by *p*_res_(*t*), a unit amplitude pulse that is active for a duration *t*_res_ after a spike (declared when *v*_s_ ≥ 10), to model the neuron’s refractory period. *g*_K_ models a high-threshold potassium conductance and is only activated when a spike occurs, decaying afterwards. We modeled its dynamics as

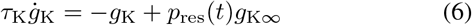

where *τ*_K_ is the decay time constant.

The dendrite compartment’s dimensionless model is

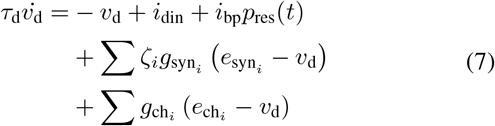

where *v*_d_ is the membrane potential, *τ*_d_ the membrane time constant, *i*_din_ the input current, *g*_syn_i__, *e*_syn_i__ and *ζ_i_* the conductance, reversal potential and decay factor, respectively, of synapse *i*, and *g*_ch_i__ and *e*_ch_i__ the conductance and reversal potential, respectively, of ion-channel population *i*. A current *i*_bp_ is injected for a duration *t*_res_ (same as in equation (5)) to model a backpropagating spike.

The decay factor, which models the spatial decay in dendritic trees, is given by^4^

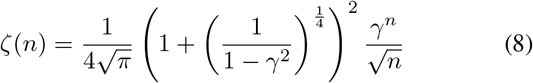

where *γ* is the silicon dendritic tree’s decay constant and *n* the distance traveled in number of neurons.

The synaptic population’s dimensionless model^5^ is

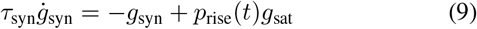

where *g*_syn_ is the instantaneous synaptic conductance, *τ*_syn_ the synaptic time constant and *g*_sat_ the maximum conductance. The unit amplitude pulse *p*_rise_(*t*)’s width *t*_rise_ models the duration for which neurotransmitter is available in the cleft following a pre-synaptic spike.

The gating variable’s model is:

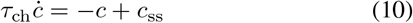

where *c*_ss_ is its steady-state activation (or inactivation) and *τ*_ch_ its time constant. *c*_ss_ is given by

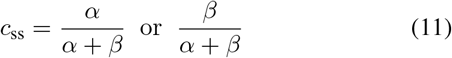

where *α* and *β* model a channel’s opening and closing rates and are descibed by

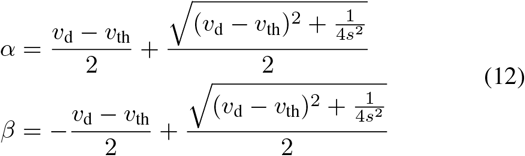

Here, *v*_th_ is the membrane potential at which *c*_s_ = 1/2 and *s* is the slope at this point. *α* and *β* satisfy a difference relation, *α* – *β* = *v*_d_ – *v*_th_, and a reciprocal relation, *αβ* = 1/ (16*s*^2^), resulting in a sigmoidal dependence of *c*_ss_ on *v*_d_. The gating variable’s time constant is given by

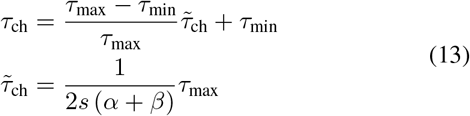

*τ*_ch_ is bell-shaped with a maximum value of *τ*_max_ when *v*_d_ = *v*_th_ and a minimum value of *τ*_min_ when |*v*_d_ – *v*_th_ | » 1/(2*s*), to avoid unphysiologically short time constants.

One or two gating variables may be used to model a channel population. When using one gating variable, its associated maximum conductance *g*_max_ may be set by a synapse population’s *g*_syn_ to model an ion channel population that is voltage-as well as ligand-gated (e.g., NMDA receptors). When using a pair of gating variables, the effective conductance is given by:

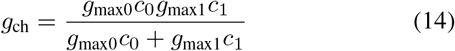

where *g*_max0_ and *g*_max1_ are the maximum conductances associated with gating variables *c*_0_ and *c*_1_, respectively. In this case, *g*_max0_ and *g*_max1_ may be set by two synapse populations’ *g*_syn_ to model neuromodulation; the gating variables’ thresholds are set low to eliminate voltage-dependence. Alternatively, *g*_max0_ and *g*_max1_ may be programmed to the same value, *g*_max_, to model a channel that activates and inactivates. In this case, the above expression simplifies to 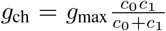. The pair of gating variables always share a programmable reversal potential *e*_ch_.

The parameter values used to obtain the simulation results presented in Figs 2 to 4 are listed in Tables 1 to 4. Reversal potentials and gating-variable thresholds were converted to dimensionless form by assuming that the leak’s reversal potential is −80 mV and the spike threshold is −50 mV (i.e., all voltages are normalized by *V*_th_ = 30 mV).

**Table 1.** Parameter values for models of cortical neuron types. Current injection for FS, RS and CH neuron types was modeled with *i*_sin_ and for IB neuron type with *i*_din_. Parameter values were chosen to produce the best fit to the *in vitro* data. All time values are in ms.

|  |  |  |  |  |  |  |  |  | Channel |  |  |  |  |  |  |  |  |
| --- | --- | --- | --- | --- | --- | --- | --- | --- | --- | --- | --- | --- | --- | --- | --- | --- | --- |
|  | Soma |  |  |  |  | Dendrite |  |  | Conductance |  | Activation |  |  | Inactivation |  |  |  |
| | $\tau_s$ | $\tau_K$ | $g_{K\infty}$ | $t_{\text{res}}$ | $i_{\text{sin}}$ | $\tau_d$ | $i_{\text{bp}}$ | $i_{\text{din}}$ | $g_{\text{max}}$ | $e_{\text{ch}}$ | $\tau_{\text{max}}$ | $v_{\text{th}}$ | $s$ | $\tau_{\text{max}}$ | $v_{\text{th}}$ | $s$ | |
| FS | 3 | 200 | 0.005 | 0.8 | 3.7-9.8 | - | - | - | - | - | - | - | - | - | - | - |  |
| RS | 15 | 200 | 50 | 0.1 | 0.08-1.42 | - | - | - | - | - | - | - | - | - | - | - |  |
| IB | 18 | 200 | 50 | 1 | 0 | 54 | 0 | 1.09-2.5 | 1 | 7.5 | 1 | 0.5 | 1.25 | 50 | 0.2 | -0.5 |  |
| CH | 13 | 50 | 250 | 2 | 30-39 | 12 | 100 | 0 | - | - | - | - | - | - | - | - |  |

**Table 2.** Parameter values used to model NMDA spikes in basal dendrites. NMDA and AMPA reversal potentials as well as NMDA threshold were chosen to match physiological values^6^. Other parameter values were chosen to produce the best fit to the *in vitro* data. All time values are in ms.

| Soma |  |  |  |  | Dendrite |  |  | NMDA |  |  |  |  |  | AMPA |  |  |  |
| --- | --- | --- | --- | --- | --- | --- | --- | --- | --- | --- | --- | --- | --- | --- | --- | --- | --- |
| $\tau_s$ | $\tau_K$ | $g_{K\infty}$ | $t_{\text{res}}$ | $i_{\text{sin}}$ | $\tau_d$ | $i_{\text{din}}$ | $i_{\text{bp}}$ | $\tau_{\text{syn}}$ | $t_{\text{rise}}$ | $g_{\text{sat}}$ | $e_{\text{syn}}$ | $v_{\text{th}}$ | $s$ | $\tau_{\text{syn}}$ | $t_{\text{rise}}$ | $g_{\text{sat}}$ | $e_{\text{syn}}$ |
| 20 | - | - | 1 | 0 | 30 | 0 | 0 | 150 | 4 | 500 | 2.7 | 2.3 | 1 | 7.25 | 0.6 | 25-240 | 2.7 |

**Table 3.** Parameter values used to model coincidence detection in apical dendrites. Soma compartment was modeled with RS neuron parameter values (see Table 1). Sodium channel parameters are similar to Mainen et. al.^7^, adjusted together with other parameter values to produce the best fit to the *in vitro* data. All time values are in ms.

|  |  |  |  |  |  |  |  | Sodium channel |  |  |  |  |  |  |  |
| --- | --- | --- | --- | --- | --- | --- | --- | --- | --- | --- | --- | --- | --- | --- | --- |
| Dendrite |  |  | Synapse |  |  |  |  | Conductance |  | Activation |  |  | Inactivation |  |  |
| $\tau_d$ | $i_{\text{din}}$ | $i_{\text{bp}}$ | $\tau_{\text{syn}}$ | $t_{\text{rise}}$ | $g_{\text{sat}}$ | $e_{\text{syn}}$ | | $g_{\max}$ | $e_{\text{ch}}$ | $\tau_{\max}$ | $v_{\text{th}}$ | $s$ | $\tau_{\max}$ | $v_{\text{th}}$ | $s$ |
| 3 | 0 | 100 | 8 | 4 | 80 | 2.7 |  | 10 | 5 | 0.4 | 0.9 | 1.2 | 35 | 0.5 | -2 |

**Table 4.** Parameter values for attentional modulation simulation. Excitatory and inhibitory neurons’ soma compartments were modeled with RS and FS neuron parameter values, respectively (see Table 1). NMDA, AMPA and dendrite were modeled with the parameter values for NMDA spikes in basal dendrites (see Table 2). All time values are in ms.

| FEF Excitatory |  |  |  |  |  |  |  |  |  |  |  |  |  |  |  |
| --- | --- | --- | --- | --- | --- | --- | --- | --- | --- | --- | --- | --- | --- | --- | --- |
| Soma |  |  |  |  | AMPA |  |  |  |  | GABA |  |  |  |  |  |
| $\tau_s$ | $\tau_K$ | $g_{K\infty}$ | $t_{\text{res}}$ | $i_{\text{sin}}$ | $\tau_{\text{syn}}$ | $t_{\text{rise}}$ | $g_{\text{sat}}$ | $e_{\text{syn}}$ | $\gamma$ | $\tau_{\text{syn}}$ | $t_{\text{rise}}$ | $g_{\text{sat}}$ | $e_{\text{syn}}$ | $\gamma$ | |
| 15 | 200 | 50.0 | 0.1 | 0 | 7.25 | 0.6 | 8 | 2.7 | 0.65 | 15 | 1 | 0.01 | 0.1 | 0.97 |  |

| FEF Inhibitory |  |  |  |  |  |  |  |  |  |  |
| --- | --- | --- | --- | --- | --- | --- | --- | --- | --- | --- |
| Soma |  |  |  |  | AMPA |  |  |  |  |  |
| $\tau_s$ | $\tau_K$ | $g_{K\infty}$ | $t_{\text{res}}$ | $i_{\text{sin}}$ | $\tau_{\text{syn}}$ | $t_{\text{rise}}$ | $g_{\text{sat}}$ | $e_{\text{syn}}$ | $\gamma$ | |
| 3 | 200 | 0.005 | 0.8 | 0.15 | 7.25 | 0.6 | 10 | 2.7 | 0.6 |  |

**Table 4 | Parameter values for attentional modulation simulation.**
| V4 Excitatory |  |  |  |  |  |  |  |  |  |  |  |  |  |  |  |  |  |
| --- | --- | --- | --- | --- | --- | --- | --- | --- | --- | --- | --- | --- | --- | --- | --- | --- | --- |
| Soma |  |  |  |  | Dendrite |  | AMPA |  |  |  |  | NMDA |  |  |  |  |  |
| $\tau_s$ | $\tau_K$ | $g_{K\infty}$ | $t_{\text{res}}$ | $i_{\text{sin}}$ | $\tau_d$ | $i_{\text{din}}$ | $\tau_{\text{syn}}$ | $t_{\text{rise}}$ | $g_{\text{sat}}$ | $e_{\text{syn}}$ | $\gamma$ | $\tau_{\text{syn}}$ | $t_{\text{rise}}$ | $g_{\text{sat}}$ | $e_{\text{syn}}$ | $v_{\text{th}}$ | $s$ |
| 15 | 200 | 50 | 0.1 | 0 | 30 | 0.001 | 7.25 | 0.6 | 2.0 | 2.7 | 0.65 | 150 | 4 | 50 | 2.7 | 2.3 | 1 |

### Programming Neurogrid

Neuronal and synaptic model parameters and connectivity are described in Python, similar to PyNN^8^, and a C++ back-end uses this description to program Neurogrid’s electronic circuit parameters and lookup tables. Examples of Python descriptions of soma, dendrite, synapse and gating variables are listed in Table 5. A calibration procedure is used to obtain a set of conversion factors to map the neuronal and synaptic model parameters to the circuit parameters^9^. In a population of silicon neurons, these parameters are lognormally distributed with a coefficient of variation (CV) of about 10% (e.g., 1/*τ*_s_ has a CV of 7.2%). Lookup tables for intracolumn connections are stored in the Neuorocores’ 256 × 16-bit RAM and for intercolumn connections in the daughterboard’s 8M × 32-bit RAM.

**Table 5.**
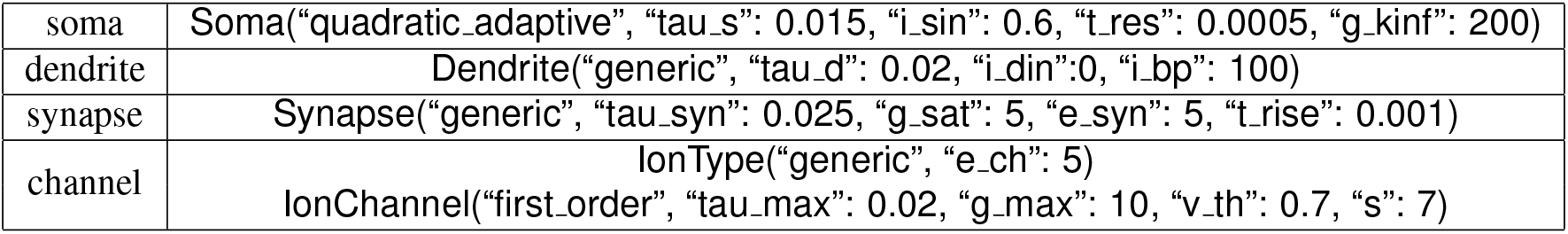
Python constructs to program Neurogrid. The table lists example Python function calls for modeling a soma, dendrite, synapse and channel. All time values are in seconds.

## Footnotes

i (250 J/s × 510 s)/(24G syn × 25 spk/s × 1*s*) = 212n J/syn/spk

## References

1. Abbott, L. F. et al. The mind of a mouse. Cell 182, 1372–1376 (2020).

2. Urai, A. E., Doiron, B., Leifer, A. M. & Churchland, A. K. Large-scale neural recordings call for new insights to link brain and behavior. arXiv preprint arXiv:2103.14662 (2021).

3. Markram, H. The Human Brain Project. Scientific American 306, 50–55 (May 2012).

4. Knight, J. C. & Nowotny, T. Larger GPU-accelerated brain simulations with procedural connectivity. Nature Computational Science 1, 136–142 (2021).

5. Boahen, K. Point-to-point connectivity between neuromorphic chips using address events. Circuits and Systems II: Analog and Digital Signal Processing, IEEE Transactions on 47, 416–434 (2000).

6. Mahowald, M. An analog VLSI system for stereoscopic vision (Springer, 1994).

7. Vogelstein, R. J., Mallik, U., Culurciello, E., Cauwenberghs, G. & Etienne-Cummings, R. A Multichip Neuromorphic System for Spike-Based Visual Information Processing. Neural Computation 19, 2281–2300 (Sept. 2007).

8. Benjamin, B. V. et al. Neurogrid: A mixed-analog-digital multichip system for large-scale neural simulations. Proceedings of the IEEE 102, 699–716 (2014).

9. Chklovskii, D. B. et al. Synaptic Connectivity and Neuronal Morphology-Two Sides of the Same Coin. Neuron 43, 609–618 (2004).

10. Binzegger, T. et al. Stereotypical Bouton Clustering of Individual Neurons in Cat Primary Visual Cortex. Journal of Neuroscience 27, 12242–12254 (Nov. 2007).

11. Merolla, P., Arthur, J., Alvarez, R., Bussat, J.-M. & Boahen, K. A multicast tree router for multichip neuromorphic systems. IEEE Transactions on Circuits and Systems I: Regular Papers 61, 820–833 (2013).

12. Park, J., Yu, T., Joshi, S., Maier, C. & Cauwenberghs, G. Hierarchical address event routing for reconfigurable large-scale neuromorphic systems. IEEE transactions on neural networks and learning systems 28, 2408–2422 (2016).

13. Choi, T. Y., Merolla, P. A., Arthur, J. V., Boahen, K. A. & Shi, B. E. Neuromorphic implementation of orientation hypercolumns. IEEE Transactions on Circuits and Systems I: Regular Papers 52, 1049–1060 (2005).

14. Izhikevich, E. M. Dynamical systems in neuroscience (The MIT Press, 2007).

15. Mountcastle, V. B. Modality and topographic properties of single neurons of cat’s somatic sensory cortex. Journal of Neurophysiology 20, 408–34 (July 1957).

16. Hubel, D. H. & Wiesel, T. N. Ferrier Lecture: Functional Architecture of Macaque Monkey Visual Cortex. Proceedings of the Royal Society of London. Series B, Biological Sciences 198, 1–59 (May 1977).

17. Stettler, D. D., Das, A., Bennett, J. & Gilbert, C. D. Lateral connectivity and contextual interactions in macaque primary visual cortex. Neuron 36, 739–750 (Nov. 2002).

18. Binzegger, T., Douglas, R., Martin, K. & Binzegger, T. A Quantitative Map of the Circuit of Cat Primary Visual Cortex. Journal of Neuroscience 24, 8441–8453 (Sept. 2004).

19. Andreou, A. G. & Boahen, K. A. Translinear circuits in subthreshold MOS. Analog Integrated Circuits and Signal Processing 9, 141–166 (1996).

20. Arthur, J. V. & Boahen, K. A. Synchrony in Silicon: The Gamma Rhythm. IEEE Transactions on Neural Networks 18, 1815–1825 (Nov. 2007).

21. Gao, P., Benjamin, V. & Boahen, K. Dynamical System Guided Mapping of Quantitative Neuronal Models Onto Neuromorphic Hardware. IEEE Transactions on Circuits and Systems I: Regular Papers 59, 2383–2394 (Oct. 2012).

22. Kauderer-Abrams, E. & Boahen, K. Calibrating silicon-synapse dynamics using time-encoding and decoding machines in 2017 IEEE International Symposium on Circuits and Systems (ISCAS) (2017), 1–4.

23. Hodgkin, A. L. & Huxley, A. F. A quantitative description of membrane current and its application to conduction and excitation in nerve. The Journal of Physiology 117, 500–544 (Aug. 1952).

24. Nowak, L. G., Azouz, R., Sanchez-Vives, M. V., Gray, C. M. & McCormick, D. A. Electrophysiological classes of cat primary visual cortical neurons in vivo as revealed by quantitative analyses. Journal of Neurophysiology 89, 1541–66 (Mar. 2003).

25. Connors, B. W., Gutnick, M. J. & Prince, D. A. Electrophysiological properties of neocortical neurons in vitro. Journal of Neurophysiology 48, 1302–1320 (1982).

26. Gray, C. M. & McCormick, D. A. Chattering Cells: Superficial Pyramidal Neurons Contributing to the Generation of Synchronous Oscillations in the Visual Cortex. Science 274, 109–113 (Oct. 1996).

27. McCormick, D. A., Connors, B. W., Lighthall, J. W. & Prince, D. A. Comparative electrophysiology of pyramidal and sparsely spiny stellate neurons of the neocortex. Journal of Neurophysiology 54, 782–806 (1985).

28. Mainen, Z. F. & Sejnowski, T. J. Influence of dendritic structure on firing pattern in model neocortical neurons. Nature 382, 363–366 (1996).

29. Arthur, J. & Boahen, K. Silicon-Neuron Design: A Dynamical Systems Approach. Circuits and Systems I: Regular Papers, IEEE Transactions on 58, 1034–1043. ISSN: 1549-8328 (May 2011).

30. Schiller, J., Major, G., Koester, H. J. & Schiller, Y. NMDA spikes in basal dendrites of cortical pyramidal neurons. Nature 404, 285–288 (2000).

31. Stuart, G. J. & Hausser, M. Dendritic coincidence detection of EPSPs and action potentials. Nature Neuroscience 4, 63–71 (2001).

32. Lavzin, M., Rapoport, S., Polsky, A., Garion, L. & Schiller, J. Nonlinear dendritic processing determines angular tuning of barrel cortex neurons in vivo. Nature 490, 397–401 (Sept. 2012).

33. Xu, N.-l. et al. Nonlinear dendritic integration of sensory and motor input during an active sensing task. Nature 492, 247–251 (Nov. 2012).

34. McAdams, C. J. & Maunsell, J. H. R. Effects of attention on orientation-tuning functions of single neurons in macaque cortical area V4. The Journal of Neuroscience 19, 431–441 (1999).

35. Self, M. W., Kooijmans, R. N., Supèr, H., Lamme, V. A. & Roelfsema, P. R. Different glutamate receptors convey feedforward and recurrent processing in macaque V1. Proceedings of the National Academy of Sciences 109, 11031–11036 (2012).

36. Peyrache, A. et al. Spatiotemporal dynamics of neocortical excitation and inhibition during human sleep. Proceedings of the National Academy of Sciences 109, 1731–1736 (Jan. 2012).

37. Ardid, S., Wang, X.-J. & Compte, A. An integrated microcircuit model of attentional processing in the neocortex. The Journal of Neuroscience 27, 8486–8495 (2007).

38. Mahowald, M. & Douglas, R. A silicon neuron. Nature 354, 515–8 (Dec. 1991).

39. Van Schaik, A. Building blocks for electronic spiking neural networks. Neural networks 14, 617–628 (2001).

40. Hynna, K. M. & Boahen, K. Thermodynamically equivalent silicon models of voltage-dependent ion channels. Neural Computation 19, 327–350 (2007).

41. Wang, Y. & Liu, S.-C. A two-dimensional configurable active silicon dendritic neuron array. Circuits and Systems I: Regular Papers, IEEE Transactions on 58, 2159–2171 (2011).

42. Benjamin, B. V., Arthur, J. V., Gao, P., Merolla, P. & Boahen, K. A superposable silicon synapse with programmable reversal potential. 2012 Annual International Conference of the IEEE Engineering in Medicine and Biology Society, 771–774 (Aug. 2012).

43. Ramakrishnan, S., Wunderlich, R., Hasler, J. & George, S. Neuron array with plastic synapses and programmable dendrites. IEEE transactions on biomedical circuits and systems 7, 631–642 (2013).

44. Chicca, E., Stefanini, F., Bartolozzi, C. & Indiveri, G. Neuromorphic electronic circuits for building autonomous cognitive systems. Proceedings of the IEEE 102, 1367–1388 (2014).

45. Schemmel, J., Kriener, L., Müller, P. & Meier, K. An accelerated analog neuromorphic hardware system emulating NMDA-and calcium-based nonlinear dendrites in 2017 International Joint Conference on Neural Networks (IJCNN) (2017), 2217–2226.

## References

1. Knight, B. W. Dynamics of encoding in a population of neurons. The Journal of General Physiology 59, 734–766 (1972).

2. Ermentrout, G. B. & Kopell, N. Parabolic bursting in an excitable system coupled with a slow oscillation. SIAM Journal on Applied Mathematics 46, 233–253 (1986).

3. Latham, P. E., Richmond, B. J., Nelson, P. G. & Nirenberg, S. Intrinsic Dynamics in Neuronal Networks. I. Theory. Journal of Neurophysiology 83, 808–827 (2000).

4. Feinstein, D. I. The hexagonal resistive network and the circular approximation. Technical Report. CaltechCSTR:1988.cs-tr-88-07 (1988).

5. Destexhe, A., Mainen, Z. F. & Sejnowski, T. J. An efficient method for computing synaptic conductances based on a kinetic model of receptor binding. Neural Computation 6, 14–18 (Jan. 1994).

6. Jahr, C. E. & Stevens, C. F. Voltage dependence of NMDA-activated macroscopic conductances predicted by single-channel kinetics. The Journal of Neuroscience 10, 3178–3182 (1990).

7. Mainen, Z. F., Joerges, J., Huguenard, J. R. & Sejnowski, T. J. A model of spike initiation in neocortical pyramidal neurons. Neuron 15, 1427–1439. ISSN: 0896-6273 (1995).

8. Davison, A. P. et al. PyNN: A Common Interface for Neuronal Network Simulators. Frontiers in Neuroinformatics 2, 1–10 (Jan. 2008).

9. Gao, P., Benjamin, V. & Boahen, K. Dynamical System Guided Mapping of Quantitative Neuronal Models Onto Neuromorphic Hardware. IEEE Transactions on Circuits and Systems I: Regular Papers 59, 2383–2394 (Oct. 2021).

